# Protein phosphatase 4 is required for Centrobin function in DNA damage repair

**DOI:** 10.1101/2023.05.31.542826

**Authors:** Zsuzsánna Réthi-Nagy, Edit Ábrahám, Rita Sinka, Szilvia Juhász, Zoltán Lipinszki

**Affiliations:** Biological Research Centre, Institute of Biochemistry, MTA SZBK Lendület Laboratory of Cell Cycle Regulation, ELKH, Szeged, Hungary; Doctoral School of Biology, Faculty of Science and Informatics, University of Szeged, Szeged, Hungary; Biological Research Centre, Institute of Genetics, National Biotechnology Laboratory, Szeged, Hungary; Department of Genetics, University of Szeged, Szeged, Hungary; Biological Research Centre, Institute of Biochemistry, ELKH, Szeged, Hungary

**Keywords:** Protein Phosphatase 4, Centrobin, DNA damage response, Homologous recombination repair, arms-closed chromosome morphology, DNA double-stranded break

## Abstract

Genome stability in human cells relies on the efficient repair of double-stranded DNA breaks, which is mainly achieved by homologous recombination (HR). Among the regulators of various cellular functions, Protein Phosphatase 4 (PP4) plays a pivotal role in coordinating the cellular response to DNA damage. Meanwhile, Centrobin (Ctb), initially recognized for its association with centrosomal function and microtubule dynamics, has sparked interest due to its potential contribution to DNA repair processes. In this study, we investigate the involvement of PP4 and its interaction with Ctb in HR-mediated DNA repair in human cells. Employing a range of experimental strategies, we investigate the physical interaction between PP4 and Ctb and shed light on the importance of two specific motifs in Ctb, the PP4-binding FRVP and the ATR kinase recognition SQ sequences, in the DNA repair process. Moreover, we examine cells lacking PP4 or Ctb and cells harboring FRVP and SQ mutations in Ctb, which result in similarly abnormal chromosome morphologies. This phenomenon likely results from the impaired resolution of Holliday junctions, which serve as crucial intermediates in HR. Taken together, our results provide new insights into the intricate mechanisms and interrelationships of PP4 and Ctb in the regulation of HR repair.

## Introduction

The DNA damage response (DDR) is a surveillance system evolved to safeguard genome integrity in the face of genotoxic stress throughout an organism’s life. Upon DNA damage, a complex signaling pathway activates the DDR to coordinate the detection of DNA lesions (checkpoint activation), cell cycle arrest, and efficient DNA repair (Waterman et al., 2020). Following successful repair, DDR activates checkpoint recovery to resume cell cycle progression (Smits et al., 2020; van Vugt et al., 2005). One of the most severe forms of DNA damage is double-stranded breaks (DSBs), which require immediate repair through processes such as homologous recombination (HR) or non-homologous end joining (NHEJ) (Vitor et al., 2020). Dysregulation of these processes may produce gene mutations, chromosome aberration and aneuploidy (Wechsler et al., 2011), hallmarks of proliferative diseases and genetic disorders (Jackson and Bartek, 2009; Jeggo et al., 2016). Therefore, precise control and fidelity are crucial during the repair process.

Reversible protein phosphorylation is a highly conserved molecular mechanism in eukaryotes that orchestrates complex cellular processes, including cell cycle progression and DNA damage response (Moura and Conde, 2019; Sancar et al., 2004). Phosphorylation is catalyzed by protein kinases and reversed by protein phosphatases, and serves as a molecular switch, modulating the activity, structure, half-life, localization and interacting partners of target proteins. Checkpoint activation and DNA repair are mainly regulated by the extensively studied master kinases, the Ataxia telangiectasia and Rad3-related (ATR), Ataxia-telangiectasia mutated (ATM) and DNA-dependent protein kinase (DNA-PK) (Schlam-Babayov et al., 2021), as well as their effectors, Checkpoint kinase 1 (Chk1) and Checkpoint kinase 2/Radiation sensitive 53 (Chk2/Rad53, mammal/yeast nomenclature) (Smith et al., 2020). However, dynamic phosphorylation and dephosphorylation of DDR regulators and executor molecules are critical for successful DNA repair and maintenance of genome integrity. It has become evident over the past years, that several protein phosphatases that counteract the activity of DDR kinases and dephosphorylate their substrates are involved in this process. The four major phosphatases that contribute directly to DDR regulation are the Ser/Thr Phosphoprotein phosphatase 1 (PPP1), Phosphoprotein phosphatase 2A (PPP2A) and Phosphoprotein phosphatase 4 (PPP4, hereafter PP4), as well as Cdc14, which is a dual specificity enzyme (Campos and Clemente-Blanco, 2020; Ramos et al., 2019).

PP4 is a ubiquitous and essential phosphatase that regulates a variety of cellular processes, including centrosome maturation and microtubule dynamics, cell division, development and differentiation, as well as DDR (reviewed in (Cohen et al., 2005; Park and Lee, 2020)). It has been reported that PP4 dephosphorylates several key proteins during DDR, such as ATR kinase, Rad53/Chk2 kinase, γH2AX histone, Replication protein 2A (RPA2), p53-binding protein 1 (53BP1), Deleted in breast cancer 1 protein (DBC1), and Kinesin-associated protein-1 (KAP-1) (Chowdhury et al., 2008; Hustedt et al., 2015; Lee et al., 2014; Lee et al., 2012; Lee et al., 2010a; Lee et al., 2015; Villoria et al., 2019). These findings have shed light on the necessity of PP4 in all stages of DDR, including checkpoint activation and double-stranded DNA break repair by HR, as well as NHEJ. PP4 is believed to function primarily by promoting cell recovery and the resumption of the cell cycle after successful repair (Campos and Clemente-Blanco, 2020; Park and Lee, 2020; Ramos et al., 2019).

The major form of PP4, which is present from yeast to humans, is the heterotrimeric complex consisting of an evolutionarily conserved catalytic subunit (PP4c), a scaffolding subunit (PPP4R2) and a regulatory subunit (PPP4R3, hereafter R3) (Cohen et al., 2005; Gingras et al., 2005; Lipinszki et al., 2015). The R3 subunit (Psy2 in yeast, Falafel in *Drosophila,* and PPP4R3A (hereafter R3A) and PPP4R3B (hereafter R3B) isoforms in mammals) is responsible for subcellular localization, as well as substrate recognition of PP4 (Karman et al., 2020; Lee et al., 2014; Lipinszki et al., 2015; Lyu et al., 2013; Ma et al., 2014; Sousa-Nunes et al., 2009; Torras-Llort et al., 2020; Ueki et al., 2019; Wolff et al., 2006). We and others have shown that the R3 subunit, through its non-canonical amino-terminal EVH1 domain, specifically binds to conserved short linear motifs (SLiMs: FxxP or MxPP, where x can be any amino acid) in the target proteins (Karman et al., 2020; Lipinszki et al., 2015; Ueki et al., 2019). Substitution of phenylalanine (F) and proline (P) in FxxP, or methionine and both prolines (PP) in MxPP to alanine (i.e. AxxA or AxAA, respectively), completely abolished the interaction between PP4 and its substrates (Karman et al., 2020; Ueki et al., 2019). Interestingly, these studies in *Drosophila* and human cells have identified novel FxxP/MxPP-containing DDR targets of PP4, including the Centrosomal BRCA2-interacting protein (Centrobin) (Karman et al., 2020; Ueki et al., 2019).

Centrobin (Ctb) was initially identified as a centrosomal protein that interacts with BRCA2 in human cells (Zou et al., 2005). It plays a vital role in daughter centriole duplication and elongation (Gudi et al., 2011; Zou et al., 2005), and contributes to microtubule stability and dynamics, regulated by NEK2 and Polo-like kinase 1 (Jeffery et al., 2010; Jeong et al., 2007; Lee et al., 2010b; Park and Rhee, 2013; Shin et al., 2015). Ctb has been identified as a putative substrate of ATM/ATR kinases, which recognize and phosphorylate specific motifs, like serine-glutamine (SQ) or threonine-glutamine (TQ) (Matsuoka et al., 2007). Recent studies have demonstrated that ATR kinase specifically phosphorylates Ctb in response to UV-induced DNA damage, leading to its enrichment in the nuclear matrix. This modification is essential for cell survival and proper DNA repair through HR (Ryu and Kim, 2019). However, the identity of the phosphatase responsible for dephosphorylating Ctb in DDR remains unknown. We noticed that the previously identified PP4-binding motif in human Ctb (771-FRVP-774 (Ueki et al., 2019)) is in close proximity to the putative ATR-targeted SQ motif of Ctb (781-SQ-782, Figure 1A). In addition, it is known that PP4 interacts with the ATR-interacting protein, ATRIP and, together with ATR, they co-regulate the phospho-status of different substrates (Hustedt et al., 2015). This leads to the speculation that PP4 may play a role in controlling the function of Ctb in DNA repair.

**Figure 1.**
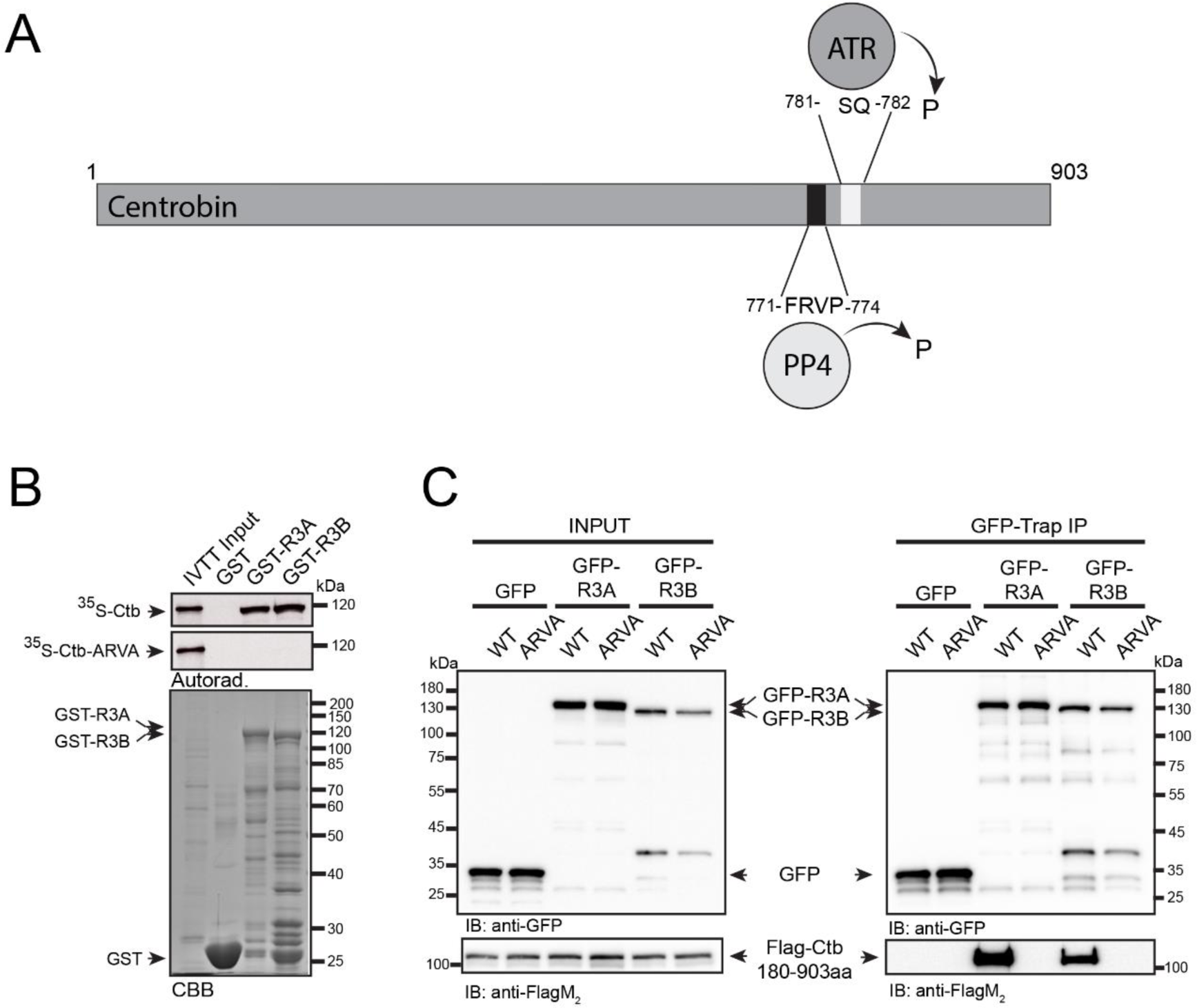
R3A and R3B bind directly to Ctb through the PP4-binding motif. **A)**. Schematic representation of the human Ctb protein: the PP4-binding FRVP sequence and the ATR kinase-recognition SQ motif are in proximity. Numbers indicate amino acid positions. P indicates putative (de)phosphorylation. **B)**. Autoradiogram (Autorad.) of GST-IVTT *in vitro* binding assay demonstrates that GST-R3A and GST-R3B (baits) specifically bind to ^35^S-methionine-labeled Ctb (^35^S-Ctb) produced in IVTT. The interaction requires an intact FRVP motif in Ctb, because the ARVA mutant (^35^S-Ctb-ARVA) does not bind to the bait proteins. The Coomassie brilliant blue-stained gel (CBB) represents the loading of the bait proteins. GST serves as a negative control bait. **C)**. GFP-Trap co-immunoprecipitation of GFP (negative control), GFP-R3A or GFP-R3A transiently co-expressed with either Flag-Ctb^180-903aa^ (indicated as WT) or Flag-Ctb^180-903aa^-ARVA (indicated as ARVA) in HEK293 cultured cells show that Ctb binding to R3A or R3B requires the intact FRVP motif *in vivo*. Cell lysate inputs and purified proteins were subjected to SDS-PAGE and western blot analysis using the indicated antibodies. Ponceau S-stained membranes are shown in Supplementary Figure 4.

In this study, we provide evidence for the physical interaction between PP4 and Ctb. We also show that the integrity of the FRVP and SQ motifs in Ctb are crucial for repairing irradiation-induced double-stranded DNA breaks through HR in human cells. Exclusion of PP4 from Ctb or mutation of the SQ motif results in a significant delay in HR and the accumulation of chromosomes with abnormal morphology, likely due to defective Holliday junction resolution (Wechsler et al., 2011; Wu et al., 2018). Our findings imply a novel function of PP4 in the late stages of HR, and further support the role of PP4 in facilitating cell cycle resumption following DNA repair.

## Results

### 1. The R3 subunit of PP4 directly binds to Ctb through its FRVP motif

It has been reported that the carboxy-terminal half of human Ctb (Ctb^460-903aa^), which contains a genuine PP4-binding motif (771-FRVP-774, Figure 1A), specifically interacts with both isoforms of R3 in HeLa cells (Ueki et al., 2019). Therefore, it was critical to understand whether the binding of full-length Ctb to R3A or R3B depends exclusively on this single FRVP motif or if additional sequences are also needed for the interaction. To test this, we performed an *in vitro* pull-down assay and found that immobilized recombinant GST-R3A and GST-R3B specifically bind to the ^35^S-methionine-labeled wild-type Ctb produced in *in vitro* coupled transcription and translation reactions (IVTT). However, when we mutated phenylalanine and proline to alanine in the FRVP motif (**<u>F</u>**RV**<u>P</u>** to **<u>A</u>**RV**<u>A</u>**), the interaction was completely abolished (Figure 1B). This indicates that Ctb has a single PP4-binding motif and that FRVP is necessary and sufficient for R3A/R3B recruitment.

We encountered difficulties when we wanted to validate the *in vitro* binding results with *in vivo* experiments using full-length Ctb. We (Supplementary figure 1), and others (Zou et al., 2005), have found that overexpressed Ctb forms aggregates around the nucleus and on the cytoskeleton and does not localize to the centrosome. Moreover, we observed the same when the amino-terminal (Ctb^1-460aa^) or carboxy-terminal (Ctb^460-903aa^) (Ueki et al., 2019) halves of Ctb were overexpressed separately in human cells (Supplementary figure 1). This may be due to the otherwise low levels of endogenous Ctb, which is tightly regulated, as well as due to interrupted structural elements in the truncated overexpressed proteins (Supplementary figure 1A). Therefore, based on secondary structure prediction, we designed a transgenic fragment of Ctb lacking the first 180 amino acids (hereafter Ctb^180-903aa^). When overexpressed, Ctb^180-903aa^ was soluble and showed centrosomal localization in human cells (Supplementary figure 1B-C). Therefore, we co-expressed Flag-tagged Ctb^180-903aa^ and its PP4-binding-deficient mutant, ARVA, together with GFP, GFP-R3A or GFP-R3B, respectively, in HEK293 cells. Using co-immunoprecipitation experiments, we demonstrated that both GFP-R3A and GFP-R3B bind to Ctb^180-903aa^, which requires an intact FRVP motif (Figure 1C). All these results imply that PP4 binds specifically to a single motif in Ctb^180-903aa^. In addition, we proved that Ctb^180-903aa^ is a soluble, centrosomal protein, with the potential for use in functional assays.

### 2. Ctb and PP4 co-operate in DNA damage repair

The variant of histone H2AX undergoes rapid phosphorylation at serine 139 (referred to as γH2AX (Rogakou et al., 1998)) in the microenvironment of chromatin surrounding DSBs (Burma et al., 2001). This modification facilitates the accumulation of DNA repair machinery (Celeste et al., 2003) and acts as a signal for DNA damage, by activating other DNA repair factors (Hunt et al., 2013). As mentioned earlier, PP4 is responsible for dephosphorylation of γH2AX, which is necessary to restore the normal progression of the cell cycle after DNA damage has been repaired (Chowdhury et al., 2008). To explore the link between Ctb and PP4 during DNA repair in the S-phase, we utilized EdU (5-ethynyl 2′-deoxyuridine) pulse-labeled HeLa cells and compared the dynamics of γH2AX in cells silenced for *ctb*, *pp4c*, *r3a*, *r3b*, or *ctb* and *pp4c*, respectively, after X-ray irradiation. We quantified the downregulation of the indicated genes by qPCR (Supplementary Figure 2B and Supplementary Table 1). All conditions showed a significant increase in the number of γH2AX nuclear foci compared to the control sample, indicating a synergistic connection between Ctb and PP4 function during the DNA repair process (Figure 2A and Supplementary Table 2). Sustained levels of γH2AX 8 hours after irradiation suggest a strong delay in DNA repair (Figure 2A, Supplementary Figure 2A). To validate these results, we repeated the experiment with a second set of siRNAs with different target sequences and found a similar phenotypic change in cells (Supplementary Figure 2C-D and Supplementary Table 2). Simultaneously, we quantified the downregulation of the indicated genes by qPCR, normalized to *actin* control (Supplementary Figure 2E and Supplementary Table 1).

**Figure 2.**
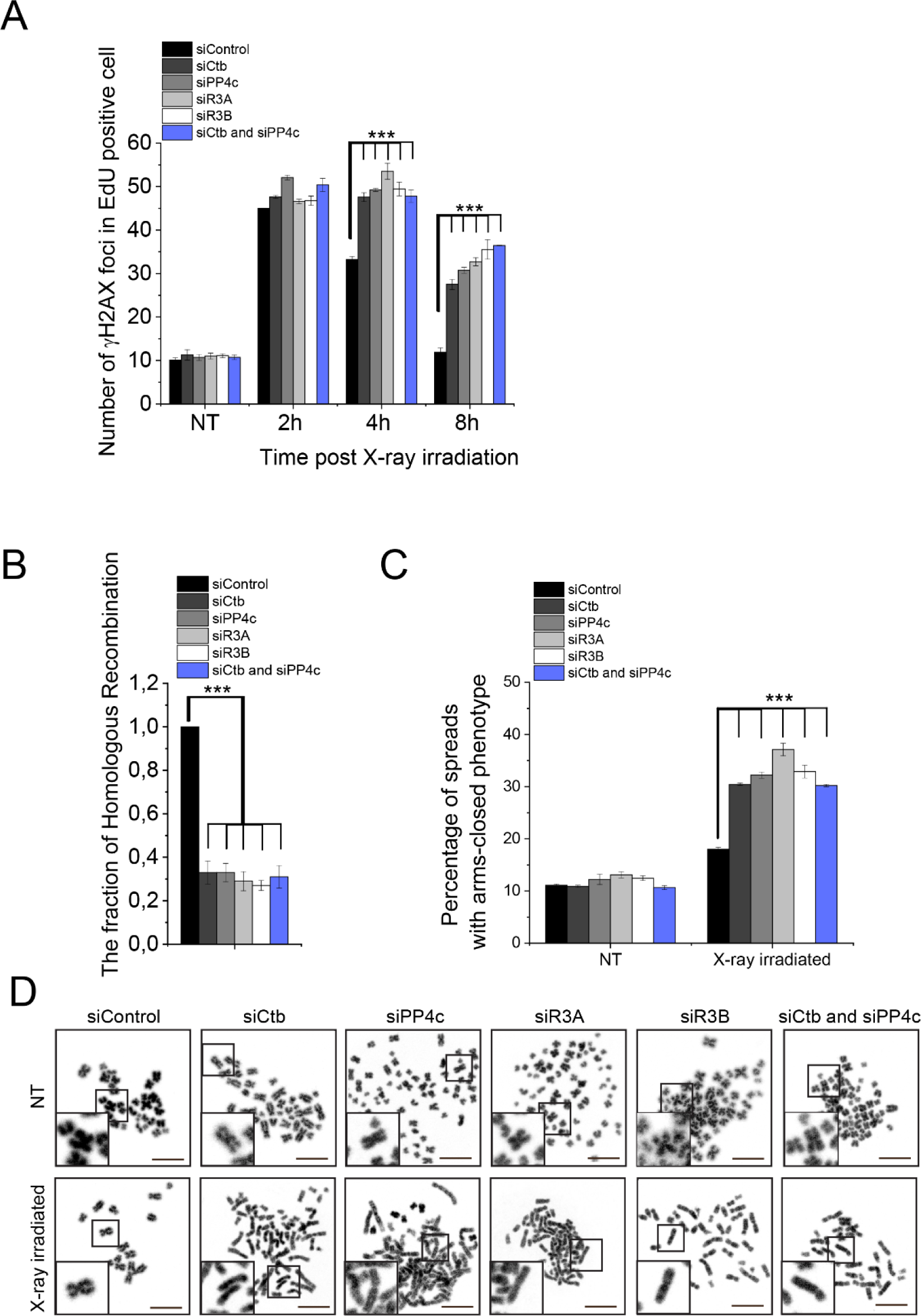
PP4 and Ctb act together during DNA damage response. **A**). Enumeration of γH2AX foci in HeLa cells. Cells were transfected with the following siRNAs: siControl, siCtb, siPP4c, siR3A, siR3B, or co-transfected with siCtb and siPP4c. After 48 h of RNAi, DNA damage was induced with 2 Gy X-ray irradiation and γH2AX foci were counted in 50 cells at different times (2, 4 and 8 h) post-irradiation. NT indicates the non-treated (non-irradiated) samples. Graphs include all data points and mean ± SEM (*n* = 3). Asterisks indicate p-values obtained by linear regression fitted independently for each time point. **B)**. Homologous recombination measuring assay. Cells were transfected with the following siRNAs: siControl, siCtb, siPP4c, siR3A, siR3B, or co-transfected with siCtb and siPP4c. After 48 h, the cells were transfected with I-SceI expressing plasmid construct. The number of GFP-positive cells was calculated with Cellprofiler (Carpenter et al., 2006), and normalized to siControl-transfected cells. Graphs include all data points and mean ± SEM (*n* = 3). Asterisks indicate p-values obtained by linear regression fitted independently for each time point. **C-D)**. Chromosome study in Ctb- or PP4 subunit-depleted cells. Cells were transfected with the following siRNAs: siControl, siCtb, siPP4c, siR3A, siR3B2, or co-transfected with siCtb and siPP4c. After 48 h, the cells were treated with 2 Gy X-ray irradiation and treated for 6 h with 20 µg/ml caffeine and 10 µg/ml colchicine. NT refers to non-treated (non-irradiated) cells. **Panel C**: Chromosome aberrations were counted in 150 spreads in each sample, and the percentage of arms closed phenotype was plotted. Graphs include all data points and mean ± SEM (*n* = 3). Asterisks indicate p-values obtained by linear regression fitted independently for each time point. **Panel D**: Representative images of chromosome spreads isolated from the indicated cells in NT or in X-ray irradiated samples. Scale bar is 10 µm. Insets at bottom left show magnified representative chromosome morphologies.

Next, we measured the frequency of HR applying the commonly used DR-GFP U2OS reporter cell line. The reporter cell line carries two truncated *gfp* (Green fluorescent protein-encoding) cassettes: the first fragment contains an I-SceI rare cutter nuclease recognition site, and the second fragment contains the missing 466 bp to prevent GFP expression by HR action. In detail, when I-SceI cuts and generates the DSB, homologous recombination takes place using the other truncated *gfp* (*igfp*) as a template to restore the GFP cassette, resulting in functional GFP. In the end, the proportion of the GFP signal-carrying cells reflects the efficiency of the HR. To apply this approach, we detected a strong decrease in HR when we downregulated *ctb*, *pp4c*, *r3a*, *r3b* separately or *ctb* and *pp4c* together (Figure 2B and Supplementary Table 2). These results confirm our previous findings and suggest that PP4 cooperates with Ctb in the regulation of HR.

To study the effect of Ctb, PP4c, R3A, R3B, or Ctb and PP4 depletions in DSB repair at the chromosome level, we isolated chromosome spreads after X-ray irradiation. We observed an increased number of chromosome aberrations, particularly a phenotype characterized by closed chromosome arms, indicating that the Holliday junction structures formed during X-ray-induced DNA repair were not properly resolved, resulting in the retention of sister chromatids. The frequency of this phenotype was significantly higher in the Ctb- or PP4-silenced cells compared to the control group (Figure 2C and 2D and Supplementary Table 2), suggesting that the depletion of these transcripts leads to a defect in DSB repair. Taken together, these results indicate that Ctb and PP4c regulate the resolution of Holliday junction structures through cooperation in HR-mediated DNA repair.

### 3. Role of the SQ and FRVP motifs of Ctb and their associated phenotypes in the DNA damage response

A SILAC (stable isotope labeling with amino acids in cell culture) study has revealed that Ser781 in the SQ motif of Ctb is phosphorylated by either ATM or ATR kinase (Matsuoka et al., 2007). Recently, it was shown that ATR kinase phosphorylates Ctb, which is required for DNA repair by HR. However, the reversing phosphatase remains unknown (Kim, 2022; Ryu and Kim, 2019). We noticed that this particular SQ motif is located adjacent to the PP4-binding site (771-FRVP-774 of Ctb, Figure 1A), which is specifically recognized by R3A and R3B subunits of PP4 (Figure 1B-C) (Ueki et al., 2019). To investigate the importance of the SQ motif (in fact, Ser781), as well as the PP4-recognition FRVP sequence in Ctb function during DNA damage response, we generated mutated forms of Ctb with the following amino acid substitutions: Ser781 in the SQ motif was replaced with alanine (S781A) to generate a non-phosphorylatable form, or with aspartate (S781D) to create a phosphor-mimetic form of Ctb; and the phenylalanine (F) and proline (P) in the FRVP motif were replaced with alanine (FRVP to ARVA) to abolish the PP4 interaction (Figure 1B-C) (Ueki et al., 2019). We first assessed the binding capacity of recombinant GST, GST-R3A, and GST-R3B to the ^35^S-labeled full-length Ctb-S781A or Ctb-S781D using the GST-IVTT *in vitro* binding assay (Rethi-Nagy et al., 2022). Interestingly, none of the mutations affected the interaction (Figure 3A). We then performed an *in vivo* co-immunoprecipitation experiment, which showed that both GFP-R3A and GFP-R3B bound to the S781A and S781D variants of Flag-Ctb^180-903aa^ (Figure 3A), similarly to the *in vitro* direct binding (Figure 3B). These results indicate that the interaction between Ctb and R3A or R3B requires the intact FRVP PP4-binding motif (Figure 1B-C), but is independent of the SQ motif of Ctb (Figure 3A-B).

**Figure 3.**
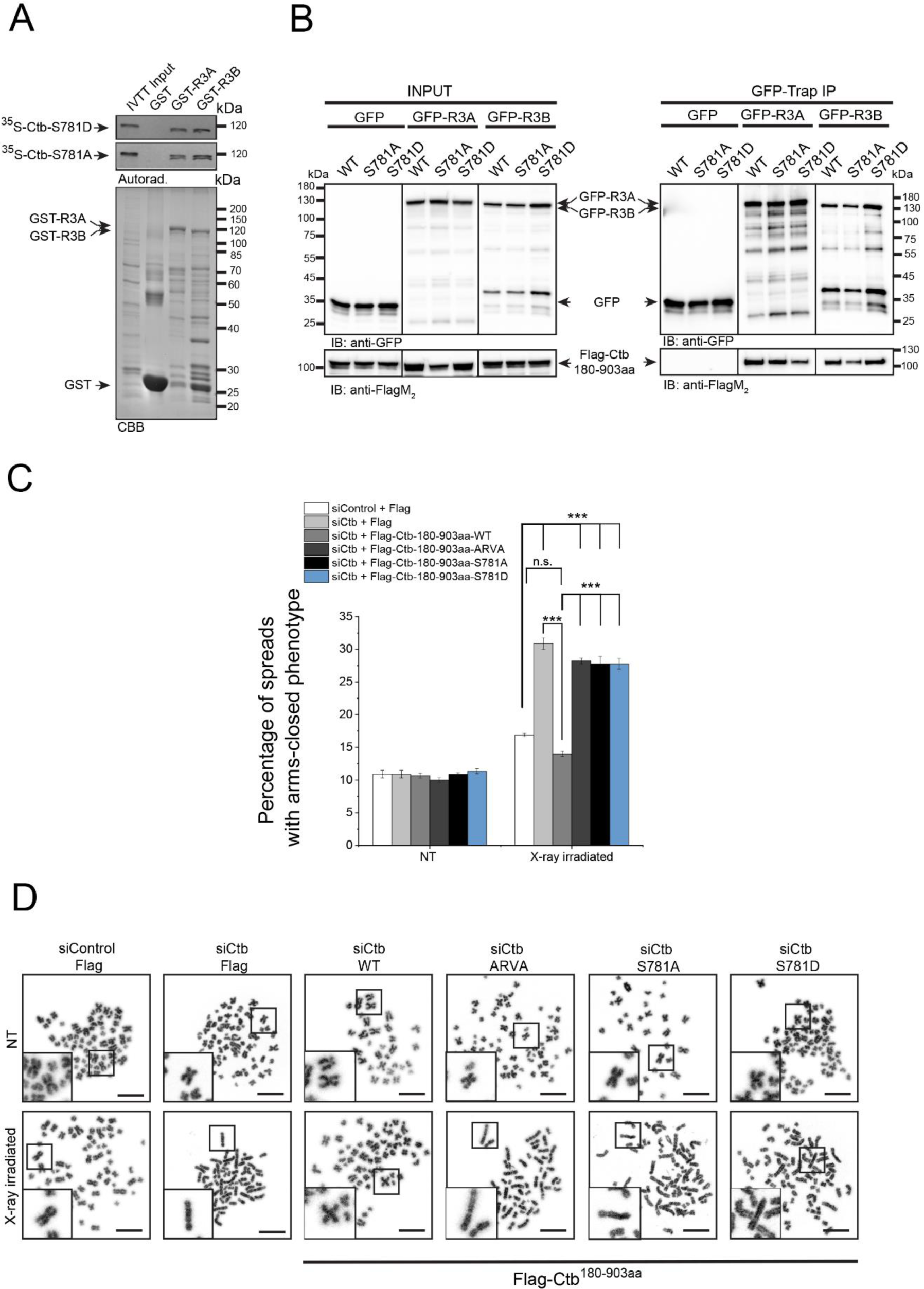
The FRVP and SQ motifs are crucial for Ctb-regulated DNA repair. **A)**. Autoradiogram (Autorad.) of GST-IVTT *in vitro* binding assay demonstrates that GST-R3A and GST-R3B (baits) interact with the S781A and S781D mutants of ^35^S-methionine-labeled Ctb (^35^S-Ctb-S781A and ^35^S-Ctb-S781D) produced in IVTT. The Coomassie brilliant blue-stained gel (CBB) represents the loading of the bait proteins. GST serves as a negative control bait. **B)**. GFP-Trap co-immunoprecipitation of GFP (negative control), GFP-R3A or GFP-R3B transiently co-expressed with either Flag-Ctb^180-903aa^ (indicated as WT), Flag-Ctb^180-903aa^-S781A (indicated as S781A) or Flag-Ctb^180-903aa^-S781D (indicated as S781D) in HEK293 cultured cells shows that Ctb binding to R3A or R3B is independent of the integrity of the SQ motif *in vivo*. Cell lysate inputs and purified proteins were subjected to SDS-PAGE and western blot analysis using the indicated antibodies. Ponceau S-stained membranes are shown in Supplementary figure 4. **C-D)**. Chromosome study in siControl or siCtb cells co-transfected with empty Flag plasmid or Ctb-depleted (siCtb) cells co-transfected with the wild type or S781A/S781D/ARVA mutant variants of Flag-Ctb^180-903aa^. After 48 h, the cells were treated with 2 Gy X-ray irradiation and treated for 6 h with 20 µg/ml caffeine and 10 µg/ml colchicine. NT refers to non-treated (non-irradiated) cells. **Panel C**: Chromosome aberrations were counted in 150 spreads in each sample, and the percentage of arms closed phenotype was plotted. Graphs include all data points and mean ± SEM (*n* = 3). Asterisks indicate p-values obtained by linear regression fitted independently for each time point. **Panel D:** Representative micrographs of chromosome spreads isolated from cells after X-ray irradiation or without treatment (NT). Scale bar is 10 µm. Insets at bottom left show magnified representative chromosome morphologies.

Next, we tested whether these Ctb variants have an impact on the modulation of DNA repair progression. We co-transfected cells with Flag-tagged wild-type Ctb^180-903aa^ or its ARVA, S781A or S781D derivatives, respectively, along with Ctb siRNA. It is important to note that the siRNA we used exclusively targeted the endogenous *ctb* transcript (its 5’ end, which is missing from the Flag-tagged mutants, Supplementary Table 1), and did not affect the levels of the transgenic proteins (Supplementary Figure 3). This allowed us to specifically assess the function of the indicated Flag-Ctb derivatives. Using this experimental setup, we performed a chromosome aberration assay and observed an increased number of abnormal closed-arm morphology (Wechsler et al., 2011) in the siCtb sample upon X-ray irradiation, which was restored in cells expressing wild-type Ctb fragment (Figure 3C and 3D and Supplementary Table 2). However, the expression of the ARVA, S781A or S781D mutation-containing Ctb derivatives did not restore the function of Ctb, suggesting that PP4 binding to Ctb, as well as the integrity of the SQ motif, is key in the regulation of DNA repair (Figure 3C and 3D).

To summarize, our results emphasize the importance of PP4 binding to the single FRVP motif, as well as the ATR-targeted SQ motif, in the functioning of Ctb, particularly in its regulation of DNA repair progression in response to DNA damage.

## Discussion

Understanding the molecular etiology of cancer plays an indispensable role in shaping the future of cancer diagnosis and personalized therapy for patients. The Ser/Thr Protein Phosphatase 4, PP4, has been extensively studied and found to play a critical role in several cellular processes, including cell cycle regulation (Archambault et al., 2022; Barr et al., 2011; Chen et al., 2007; Helps et al., 1998; Karman et al., 2020; Lipinszki et al., 2015; Park and Lee, 2020; Rocha et al., 2023; Torras-Llort et al., 2020; Toyo-oka et al., 2008; Zhuang et al., 2014) and DNA repair (Guo et al., 2022; Keogh et al., 2006; Lee et al., 2014; Lee et al., 2012; Lee et al., 2010a; O’Neill et al., 2007; Shaltiel et al., 2014). The impact of PP4 on cancer development has been investigated, revealing that alterations in PP4 expression, subcellular localization, or activity can contribute to tumorigenesis. In breast, lung, colon and prostate cancer, an increase in PP4 expression is often observed, suggesting a possible oncogenic role of this phosphatase (Li et al., 2015; Peng and Maller, 2010; Wang et al., 2008). On the other hand, a marked decrease in PP4 expression or loss of PP4 activity is observed in ovarian and cervical cancer, highlighting its tumor suppressive functions (Dong et al., 2012). Moreover, there is compelling evidence demonstrating the role of PP4 in DNA repair through its regulation and termination of signaling events (Campos and Clemente-Blanco, 2020). A specific target of PP4 is γH2AX, which becomes phosphorylated upon DNA damage. The dephosphorylation of γH2AX by PP4 is essential for restoring the normal progression of the cell cycle after the completion of DNA repair (Chowdhury et al., 2008).

Centrosomes are multifunctional regulators of genome stability and ultimately serve as scaffolds for DNA repair proteins (Lerit and Poulton, 2016). Centrosomes are the major microtubule organizing centers in animal cells and modulate the dynamics of microtubules, too, which are required for DDR protein transportation when DNA is damaged (Kim, 2022; Li et al., 2022). It has recently been shown, that the daughter centriole-specific Centrobin (Ctb), which is needed for centrosome duplication (Zou et al., 2005) and microtubule stability (regulated by the activity of NEK2 and Polo-like kinase 1) (Lee et al., 2010b; Park and Rhee, 2013; Shin et al., 2015), also localizes to DSB sites and interacts with important HR factors, indicating its direct involvement in the repair process (Ryu and Kim, 2019). Ctb has been identified as a BRCA2-interacting protein, one of the major regulators of DNA repair by HR (Zou et al., 2005). In addition, Ctb has been found to be phosphorylated by ATR/ATM kinase, further supporting its role in DNA repair (Matsuoka et al., 2007; Ryu and Kim, 2019). We, and others, have shown that Centrobin is a genuine target of PP4 (Karman et al., 2020; Ueki et al., 2019), however, whether PP4 is involved in the DDR function of Ctb has not yet been studied.

Through rigorous experiments, we have successfully confirmed a direct interaction between the regulatory subunits of PP4 phosphatase R3A or R3B, and full-length Ctb (Figure 1). We provided evidence that the interaction depends on the presence of a single PP4-binding FxxP motif (FRVP) within Ctb, with any alterations or mutations in the phenylalanine (F) and proline (P) residues potentially disrupting the binding capacity. In addition, our research has shown that knocking down Ctb, along with members of the PP4 complex (including the catalytic subunit PP4c, and the substrate-binding subunit R3A or R3B), leads to a reduction in homologous recombination frequency (Figure 2B). This suggests that both Ctb and the PP4 complex are actively involved in the same signaling pathway, highlighting their interconnectedness. Furthermore, our data suggest that Ctb, as a target of the PP4 holoenzyme, has a similar influence on the status of the histone variant H2AX, which plays a crucial role in the DNA repair response (Figure 2A). Remarkably, when we selectively silenced *ctb*, *pp4c, r3a, r3b* or the combined knockdown of *pp4c* and *ctb*, we observed a significant increase in a chromosome aberration called “arms closed” morphology (Figure 2C), the consequence of incomplete Holliday junction resolution. Furthermore, Ctb phosphorylated by ATR is required for both cell survival and homologous recombination after induction of DNA damage (Ryu and Kim, 2019). We showed that expression of ARVA (deficient in PP4-binding), and S781A or S781D (phospho-null or phospho-mimetic forms of Ser781 in the SQ motif) variants of Ctb could not rescue the Ctb-silencing induced chromosome aberration phenotype in gamma-irradiated cells (Figure 3C and 3D) compared to wild type Ctb. This finding suggests that the putative phosphorylation site within the SQ motif and the PP4 consensus recognition motif, FRVP, of Ctb are crucial in DNA repair, most probably by regulating the function of Ctb in the resolution of the Holliday structures. Based on literature data and our results, we assume that the phosphoregulation of Ctb might contribute to the interaction between Ctb and BRCA2, thereby promoting the process of homologous recombination. Notably, the precise characterization of this interaction remains the focus of future research endeavors. Considering our comprehensive results, it is evident that the PP4 holoenzyme and Ctb jointly contribute to DNA repair by homologous recombination.

In summary, our research emphasizes the interplay between PP4 and Ctb in the context of genome instability and DNA repair. By elucidating the molecular mechanisms underlying their interactions and their roles in homologous recombination, these findings support potential therapeutic targets and treatment strategies.

## Materials and methods

### 1.1. DNA constructs

cDNA encoding human PPP4R3A isoform 1 (Uniprot ID: Q6IN85; cDNA name: MHS6278-202759611, clone ID: 6142109), human PPP4R3B isoform 3 (Uniprot ID: Q5MIZ7-3, cDNA name: MHS6278-202807853; clone ID: 5259789) and human Centrobin isoform 1 (Uniprot ID: Q8N137-1; cDNA name: MHS6278-202832427, clone ID: 4859539) were originally generated by the Mammalian Gene Collection project (Gerhard et al., 2004) and we obtained from Horizon Discovery Ltd (Cambridge, UK). Coding DNA sequence (CDS) of R3A and R3B were cloned into the pDONR221 plasmid by BP reaction using the Gateway System (cat#12536017, Thermo Fisher Scientific, Waltham, MA, USA). Expression constructs were made by LR reaction using the following destination vectors: pDEST15 (for N-terminal GST-fusion in *E. coli*, cat#11802014, Thermo Fisher Scientific, Waltham, MA, USA) and pcDNA-DEST53 (for N-terminal GFP fusion in mammalian cells, cat#12288015, Thermo Fisher Scientific, Waltham, MA, USA). Modified forms of Ctb (Ctb-ARVA (771-FRVP-774 to 771-ARVA-774), S781A or S781D) were created by standard mutagenesis using QuickChange II XL Site-Directed Mutagenesis Kit (cat#200522, Agilent Technologies, Santa Clara, CA, USA). CDS of full-length Ctb, Ctb-ARVA, Ctb-S781A and Ctb-S781D were subcloned into the pHY22 plasmid (for *in vitro* expression (Rethi-Nagy et al., 2022)) by standard procedure. Truncated forms of Ctb (1-180aa, 180-903aa, 1-450aa, 450-903aa) were generated by PCR and cloned into the pFlag-CMV4 plasmid (for N-terminal Flag fusion in mammalian cells, cat#E7158, Merck Millipore, Burlington, MA, USA). CDS of ARVA, S781A and S781D mutants of Ctb^180-903aa^ were also cloned into the pFlag-CMV4 plasmid. All DNA constructs were verified by DNA sequencing. Oligonucleotide primers are provided in Supplementary Table 1.

### 1.2. Recombinant protein expression and purification

Glutathione S-transferase (GST) fused recombinant full-length R3A (GST-R3A) and R3B (GST-R3B) were expressed in Six Pack *E. coli* cells (Lipinszki et al., 2018). Bacteria cultures were grown at 16 °C for 48 h (at 280 r.p.m. using orbital shaker with 19 mm diameter) in Terrific broth autoinduction media (cat#AIMTB0210, Formedium, Hunstanton, UK), then bacteria were harvested by centrifugation (3,500 *xg*, 4 °C, 15 min) and resuspended in ice-cold phosphate-buffered saline (PBS) supplemented with 0.2 mg/mL lysozyme (cat#L6879, Sigma-Aldrich, St. Louis, MO, USA) and 1 mM phenylmethylsulfonyl fluoride (PMSF, cat#P7626, Sigma-Aldrich, St. Louis, MO, USA). Cells were lysed by standard sonication followed by centrifugation at 12,000 *xg*, 4 °C, 20 min. Recombinant proteins were affinity purified on Glutathione Sepharose 4B resin according to the manufacturer (cat#17-0756-01, Cytiva, Washington, WA, USA). Immobilized bait proteins on beads (GST, GST-R3A or GST-R3B, respectively) were stored in 50 % glycerol in PBS at −20 °C.

### 1.3. *In vitro* pull-down assay

To test the direct interaction between Ctb (prey) or its derivatives (Ctb-ARVA, Ctb-S718A and Ctb-S718D) and GST-R3A or GST-R3B (bait) protein, respectively, we performed a GST-IVTT assay as described earlier (Rethi-Nagy et al., 2022). Briefly, ^35^S-methionine-labeled Ctb or its derivatives were produced in coupled *in vitro* transcription and translation reaction (IVTT, at 30 °C for 1 h) using TNT Quick Coupled Transcription/Translation System (cat#L1170, Promega, Madison, WI, USA). Prey proteins were diluted in binding buffer (50 mM HEPES pH 7.5, 150 mM NaCl, 2 mM MgCl^2^, 1mM EGTA, 1mM DTT, 0.1% Triton X-100, EDTA-free protease inhibitor cocktail (PIC, cat#11873580001, Roche, Basel, Switzerland) and 0.5 % standard bovine serum albumin), incubated with equal amounts of immobilized GST, GST-R3A or GST-R3B proteins, respectively, and mixed at 4 °C for 2 h with gentle rotation. Beads were settled by centrifugation (500 *xg*, 4 °C, 5 min), washed three times (5 min each) with washing buffer (50 mM HEPES pH 7.5, 150 mM NaCl, 2 mM MgCl^2^, 1mM EGTA, 1mM DTT, 0.1% Triton X-100) and four times (5 min each) with washing buffer supplemented with 50 mM NaCl and 0.1% Triton X-100. Beads were boiled in Laemmli sample buffer for 4 min, IVTT inputs and eluted proteins were run on SDS-PAGE, gels were stained with Coomassie Brilliant Blue, scanned, dried, and subjected to autoradiography.

### 1.4. Cell culture maintenance

In this study all cell lines were cultured in Dulbecco’s modified Eagle’s medium (DMEM) containing GlutaMAX™ Supplement (cat#61965026, Thermo Fisher Scientific, Waltham, MA, USA) supplemented with 10% fetal bovine serum (FBS, cat#ECS0180L, Euroclone, Pero, Italy), 1xPenStrep (cat#XC-A4122, Biosera, Nuaille, France) and 1% nonessential amino acid (NEAA, cat#BE13-114E, Lonza, Basel, Switzerland) and maintained at 37 °C under 5% CO^2^, in a humidified incubator.

### 1.5. Co-immunoprecipitation

Human HEK293 cells (ATCC CRL-1573, Manassas, VA, USA) were transiently co-transfected with GFP, GFP-R3A or GFP-R3B and Flag-tagged Ctb^180-903aa^ and its ARVA, S781A or S781D mutants, respectively, using Polyethylenimine (PEI, cat#408727, Merck Millipore, Burlington, MA, USA), according to standard procedures. Cells were harvested 2 days post-transfection and lysed in Buffer A (50 mM Tris pH 7.6, 50 mM NaCl, 2 mM MgCl^2^, 0.5 mM EGTA, 0,1% NP-40, 5% glycerol, 1mM DTT, 1 mM PMSF, 1x PIC, 25 µM MG132 (cat#10012628, Cayman Chemical Company, Michigan, USA) and 0.1 µl/ml Benzonase nuclease (cat#70746-10KUN, Merck Millipore, Burlington, MA, USA) by passing the cell suspension through a G25 needle ten times. Lysates were centrifuged (17,000 *xg*, 20 min, 4 °C) and cleared supernatants were used for immunoblotting (inputs) and co-immunoprecipitations (co-IP). For co-IP clarified lysates were mixed with GFP-Trap magnetic agarose beads (cat#gtma-20, ChromoTek GmbH, Planegg, Germany) for 90 min at 4 °C, with gentle rotation. Beads were washed four times (5 min each) in Buffer B (50 mM Tris pH 7.6, 50 mM NaCl, 2 mM MgCl^2^, 0.5 mM EGTA, 0.1% NP-40, 5% glycerol), proteins were eluted by boiling in Laemmli sample buffer and analyzed by SDS-PAGE followed by immunoblotting using anti-FlagM^2^ and anti-GFP antibodies, respectively.

### 1.6. Gene silencing by RNAi and qPCR measurements of silencing efficiency

In HeLa (ATCC CRM-CCL-2, Manassas, VA, USA) or DR-GFP reporter U2OS (Gunn and Stark, 2012) cells Ctb, PP4c, R3A, and R3B were silenced separately, or Ctb and PP4c we co-silenced using the following two sets of siRNAs: Ctb (cat#4392420 siRNA ID #1: 141226, cat#AM16708 siRNA ID #2: s42056), R3A (cat#AM16708 siRNA ID #1: s31226, cat#4392420 siRNA ID #2: 133612), R3B (cat#AM16708 siRNA ID #1: s32915, cat#4392420 siRNA ID #2: 123224), PP4c (cat#AM 16708 siRNA ID #1: s10999, cat#4390824 siRNA ID #2 4441) and Control (cat#4390843). All siRNAs were purchased from Thermo Fisher Scientific (Waltham, MA, USA). Transfection of cell lines with specific siRNAs was carried out using Lipofectamine (cat#11668-018, Invitrogen, Waltham, MA, USA) or DharmaFECT^TM^ (cat#T-2022-02, GE Healthcare Dharmacon, Inc. Lafayette, Colorado, USA) transfection reagent according to the manufacturer’s protocol. Experiments were performed 48 h after siRNA transfection.

Total RNA purification was performed with Quick-RNA MiniPrep kit (cat#R1054, Zymo Research, Irvine, CA, USA). For the synthesis of first-strand cDNA the RevertAid First Strand cDNA Synthesis Kit (cat#K1670, Thermo Fisher Scientific, Waltham, MA, USA) was used according to the manufacturer’s instructions. Maxima SYBR Green/ROX qPCR Master Mix (cat#K0222, Thermo Fisher Scientific, Waltham, MA, USA) was used for the real-time quantitative PCR reaction, according to the manufacturer’s instructions. Reactions were run three times in quadruplicates in the Rotor-Gene Q Real-Time PCR Detection System (QIAGEN, Germantown, Maryland, USA) with the following reaction conditions: 95 °C 10 min, 40 cycles of 95 °C 15 sec, 55 °C 30 sec, 72 °C 30 sec. The final values represent the mean and standard error of the quadruplicates.

### 1.7. γH2AX foci quantitation assay

HeLa cells were cultivated in 6-well tissue culture plates (cat#30006, SPL Life Sciences, Pochon, Kyonggi-do, South Korea), and following 24 h of gene silencing, the cells were trypsinized (cat#TRY-3B, Capricorn Scientific, Ebsdorfergrund, Germany) using standard protocols and transferred to glass-bottom cell culture chambers (cat#631-0150, VWR International, Radnor, PA, USA). 24 h post gene silencing, the cells were exposed to 2 Gy X-rays using a Trakis XR-11 X-ray machine according to previous work (Juhasz et al., 2020). At various time points after irradiation, the cells were fixed with 3% PFA (paraformaldehyde, cat#158127, Merck Millipore, Burlington, MA, USA) for 10 min and subjected to immunofluorescence. The γH2AX foci were quantified in 50 EdU-positive S-phase cells after anti-γH2AX immunostaining. The remaining cells were collected for qPCR analysis.

### 1.8. EdU assay

For the specific labeling of S-phase cells, a medium containing 10 µM 5-ethynyl 2′-deoxyuridine (EdU, cat#BCN-001-5, BaseClick GmbH, Munich, Germany) was added to the cells 1 h before X-ray irradiation. At various time points, the cells were fixed with 3% PFA for 10 min and subjected to anti-γH2AX staining, as described below. To label the EdU positive cells, Edu-Click 555 (cat#BCK-EdU555-1, BaseClick GmbH, Munich, Germany) was used, following the manufacturer’s protocol. In the end, nuclei were counterstained with 1 μg/ml Hoechst33342 dye (cat#H21492, Thermo Fisher Scientific, Waltham, MA, USA) in PBS. Fluorescence excitation was performed using diode laser 405, 488 and 555 nm with LSM800 confocal microscope (Carl Zeiss, Jena, Germany).

### 1.9. Statistical analysis of data

All experiments were at least n=3 replicated. Statistical analysis was performed using Origin Graph statistical software (https://www.originlab.com/origin). In all cases, we made an ANOVA test (Liu and Wang, 2021). Asterisks represent *P* values, which correspond to the significance of regression coefficients (\**P* < 0.05, \*\**P* < 0.01, and \*\*\**P* < 0.001).

### 1.10. DR-GFP reporter assay

DR-GFP U2OS cells, obtained from Jeremy Stark (Gunn and Stark, 2012), were initially seeded in 6-well tissue culture plates, followed by the silencing of target genes. After 24 h, the cells were trypsinized and transferred to glass-bottom cell culture chambers. Subsequently, 24 h post-transfection, the cells were transfected with an I-SceI rare cutter endonuclease coding plasmid construct (cat #26477, Addgene Watertown, Massachusetts, USA). After 96 h of gene silencing, the cells were fixed with 3% PFA for 10 min. The cells were then visualized using confocal microscopy, and the number of GFP-positive cells, indicative of homologous recombination (HR) events, was quantified using CellProfiler software. In CellProfiler, the nuclei were segmented based on Hoechst-stained DNA. A minimum of 3000 cells were analyzed per experimental condition (Supplementary Table 2).

### 1.11. Chromosome preparation

HeLa cells were seeded in 6-well tissue culture plates and transfected with siRNAs, or co-transfected with siRNAs and Flag constructs. After 24 h, cells were separated into two 6-well plates, one was irradiated with X-ray using Trakins XR-11 X-ray machine, then treated with 20 µg/ml caffeine (cat#C0750, Merck Millipore, Burlington, MA, USA) and 10 µg/ml colchicine (cat#D00138122, Calbiochem, San Diego, CA, USA) then incubated for 6 h at 37 °C at 5% CO^2^. Cells were trypsinized and harvested with centrifugation (1000 *xg*, RT, 3 min) and incubated for 30 min at 37 °C with 75 mM KCl. Cells were washed three times with ice-cold methanol: acetic acid in a 3:1 ratio, 20 µl were dried onto coverslips and stained with Hoechst (1 μg/ml in PBS). 150 chromosome spreads were counted and classified per sample.

### 1.12. Double transfection of siRNAs and transgenic constructs

Hela cells were initially plated in a 6-well tissue culture plate and co-transfected with siControl and Flag vector, or siCtb along with Flag vector, or wild type (WT), S781A, S781D or ARVA variants of Flag-Ctb^180-903aa^. After 24 h, the cells were divided into two tissue culture plates. At 48 h post-transfection, half of the cells were subjected to X-ray treatment. Following X-ray treatment, the cells were treated with 20 µg/ml caffeine and 10 µg/ml colchicine and incubated for 6 h at 37 °C in a 5% CO^2^ environment. Chromosomes were prepared as described above.

### 1.13. Immunofluorescence and microscopy

24 h post-transfection cells were seeded onto coverslips, incubated for another 24 h followed by fixation with 3% PFA for a duration of 10 min. Subsequently, the cells were permeabilized using 0.5% Triton X-100 in PBS for 10 min, then washed three times with PBS. To block non-specific binding, the cells were incubated in a blocking buffer containing 3% bovine serum albumin and 0.1% Triton X-100 in PBS for 1 h at room temperature (RT). The samples were then treated with primary antibody diluted in blocking buffer overnight at 4 °C. After incubation with the primary antibody, the cells were washed three times with 0.1% Triton X-100 in PBS and incubated with a fluorescently tagged secondary antibody diluted in blocking buffer for 1 h at RT in the dark. Following two washes with 0.1% Triton X-100 in PBS, the cells were counterstained with Hoechst (1 μg/ml in PBS) for 10 min. Microscopy images were captured using a Zeiss LSM800 system.

### 1.14. Antibodies

For immunoblotting (IB) anti-FlagM^2^ (IB: 1:10,000, cat#F1804, Merck Millipore, Burlington, MA, USA), anti-GFP (IB: 1: 1000, cat#11814460001, Roche, Basel, Switzerland) and anti-αtubulin (IB: 1: 10,000, cat#T6199, Merck Millipore, Burlington, MA, USA) were used. For immunostaining, we used anti-γH2AX (IF: 1:500, cat#ab81299, Waltham, Boston, USA), anti-γTubulin (IF: 1:300, cat#T6557, Merck Millipore, Burlington, MA, USA) and anti-Flag (IF: 1:500, cat#F7425, Merck Millipore, Burlington, MA, USA). Secondary antibodies were the following: goat anti-mouse IgG conjugated to horseradish peroxidase (IB: 1:10,000, cat#P044701-2 Dako, Glostrup, Denmark), donkey anti-Rabbit IgG Alexa Fluor 488 (IF: 1:500, cat#A21206, Thermo Fisher Scientific, Waltham, MA, USA) and donkey anti-Mouse IgG Alexa Fluor 488 (IF: 1:500, cat#A21202, Thermo Fisher Scientific, Waltham, MA, USA), donkey anti-Rabbit IgG Alexa Fluor 594 (IF: 1:500, cat#A21203, Thermo Fisher Scientific, Waltham, MA, USA).

## Acknowledgments

The authors are grateful to Jeremy Stark (Beckman Research Institute, USA) for providing the DR-GFP U2OS cell line for the study.

## Author Contributions

Z.R.-N., S.J. and Z.L. designed the experiments and wrote the manuscript. S.J. and Z.L. supervised the project. E.A. performed co-immunoprecipitations and immunoblotting. R.S. carried out the qPCR experiments. S.J. and Z.R.-N. performed cell culture work, RNAi, immunostaining, microscopy, counting and statistical analysis. Z.R.-N. did the cloning, produced recombinant proteins, and carried out the GST-IVTT binding assay.

## Supplementary Material

Supplementary Figure 1. The localization of the various Ctb fragments in human cells.

Supplementary Figure 2. PP4 and Ctb act together during DNA damage response.

Supplementary Figure 3. The FRVP and SQ motifs are crucial for Ctb-regulated DNA repair.

Supplementary Figure 4. Uncropped images corresponding to the main and supplementary figures.

Supplementary Table 1. Oligonucleotide primers and siRNA sequences used in this study.

Supplementary Table 2. Detailed statistical analysis.

