## Supplementary Figures for "Protein phosphatase 4 is required for Centrobin function in DNA damage repair"

### Supplementary Figure 1

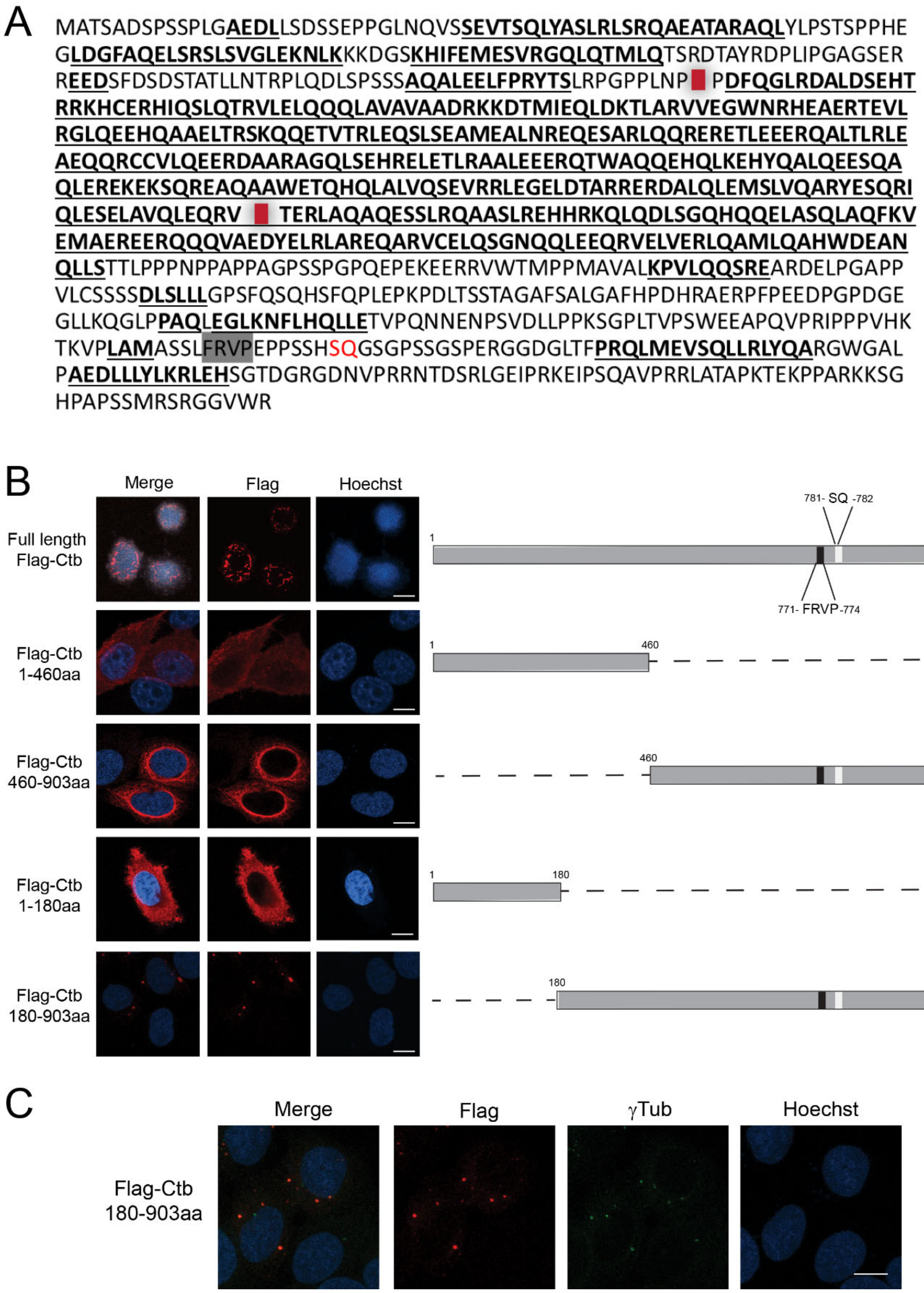


**Supplementary Figure 1. The localization of the various Ctb fragments in human cells.** **A).** The secondary structure of Ctb was predicted with PsiPred (http://bioinf.cs.ucl.ac.uk/psipred/). The endpoints of the generated fragments are indicated by red squares. Alpha-helical regions are denoted in bold and underlined. The FxxP (FRVP, highlighted in grey) motif serves as the binding site for R3 subunits, while SQ (in red) is a putative phosphorylation site targeted by ATR kinase. **B). Left panel**: Representative images of expression of full length and truncated forms of Flag-tagged Ctb. Cells were transfected with Ctb fragments-encoding plasmid construct, and 48 h post-transfection cells were fixed and stained with anti-FlagM2 (red) and Hoechst (DNA, blue). Scale bar: 10 µm. **Right panel**: The schematic representation of truncated forms of Ctb. The binding motif for R3 subunits, FRVP, and the putative phosphorylation site for ATR kinase, SQ, are indicated. Numbers indicate amino acid endpoints. **C).** Representative images of Flag-Ctb$^{180\text{-}903aa}$ fragment (red) co-localize with the centrosomal marker γ-Tub (green). Scale bar: 10 µm.

**Supplementary Figure 2**

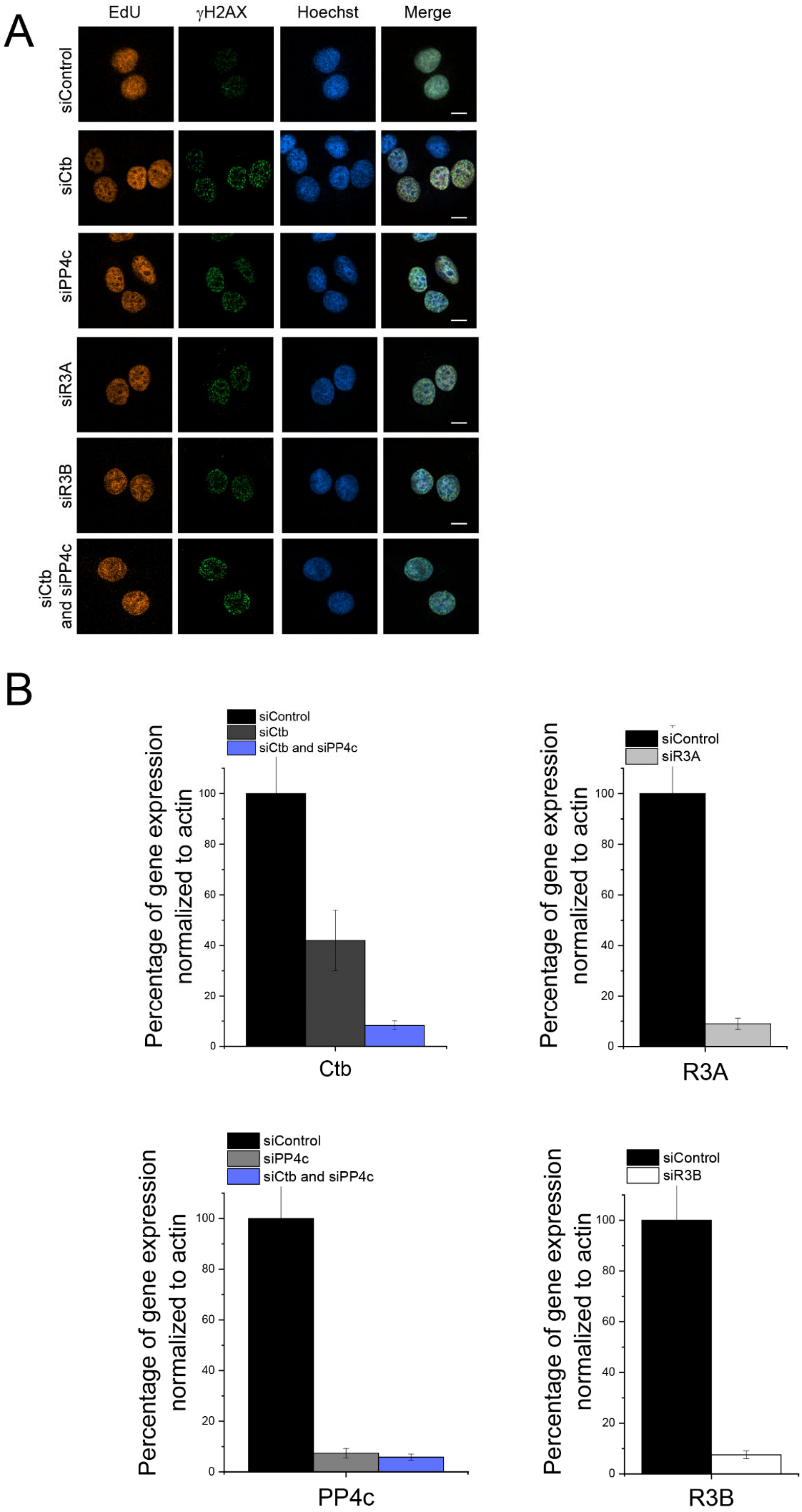

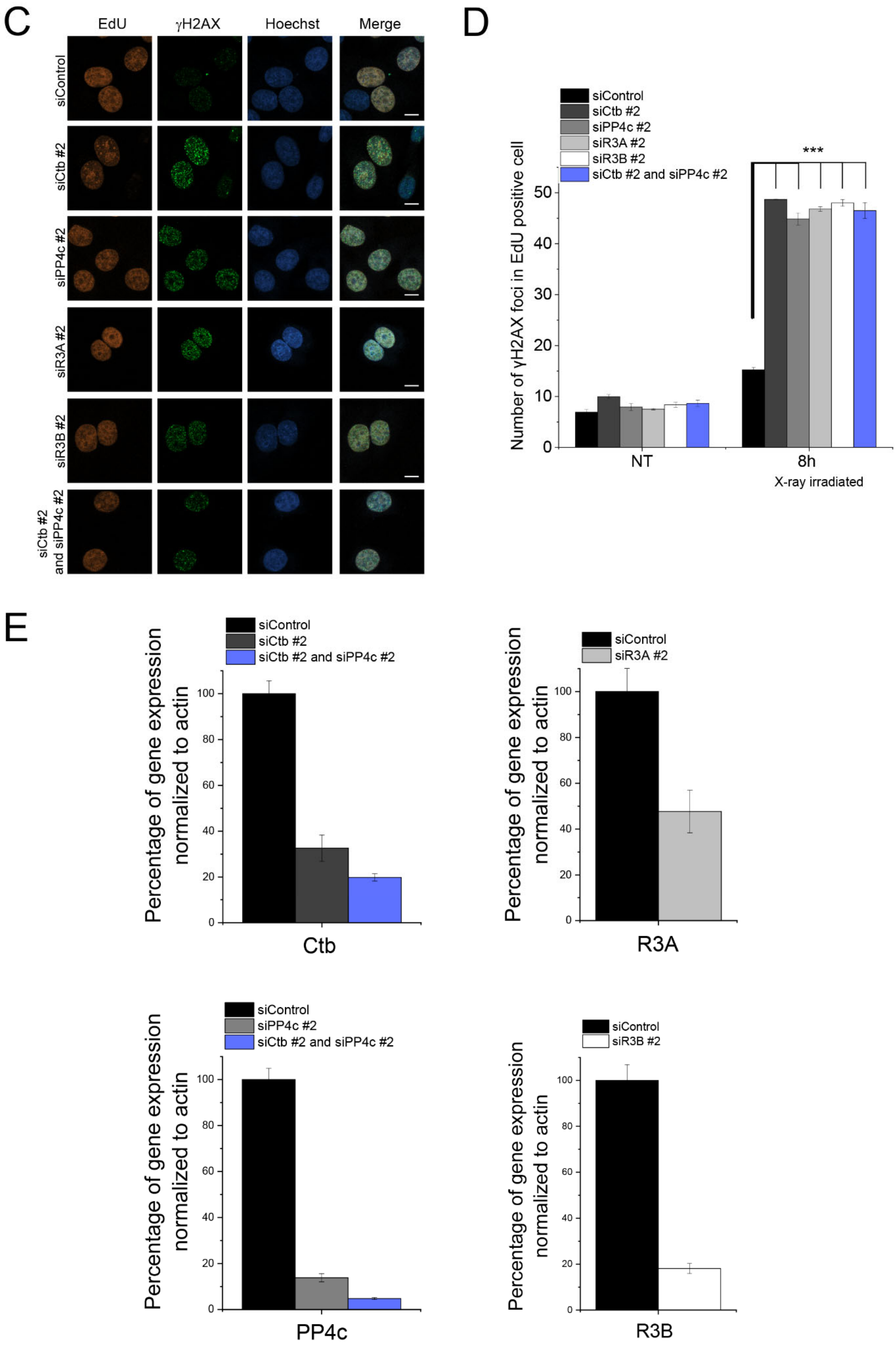

C
EdU
γH2AX
Hoechst
Merge
siControl
siCtb #2
siPP4c #2
siR3A #2
siR3B #2
siCtb #2 and siPP4c #2
D
siControl
siCtb #2
siPP4c #2
siR3A #2
siR3B #2
siCtb #2 and siPP4c #2
***
Number of γH2AX foci in EdU positive cell
0
10
20
30
40
50
NT
8h
X-ray irradiated
E
siControl
siCtb #2
siCtb #2 and siPP4c #2
Percentage of gene expression normalized to actin
0
20
40
60
80
100
Ctb
siControl
siR3A #2
Percentage of gene expression normalized to actin
0
20
40
60
80
100
R3A
siControl
siPP4c #2
siCtb #2 and siPP4c #2
Percentage of gene expression normalized to actin
0
20
40
60
80
100
PP4c
siControl
siR3B #2
Percentage of gene expression normalized to actin
0
20
40
60
80
100
R3B

**Supplementary Figure 2. PP4 and Ctb act together during DNA damage response.**

**A).** Quantification of γH2AX in HeLa cells. Cells were transfected with the following siRNAs (Set #1): siControl, siCtb, siPP4c, siR3A, or siR3B, and siCtb and siPP4c together. 48 h post-transfection the damage was induced with X-ray irradiation (2 Gy). Microscopy images represent the foci (green) 8 h after X-ray irradiation in S phase (EdU, orange) cells. Scale bar: 10 µm. **B).** Validation of gene silencing by qPCR. 48 h post-transfection cells were harvested and the level of gene expression of silenced Ctb, PP4c, R3A or R3B was measured by qPCR. The values were normalized to the level of actin. **C).** Quantification of γH2AX in HeLa cells. Cells were transfected with the second set of the following siRNAs (set #2): siControl, siCtb, siPP4c, siR3A, siR3B, and siCtb and siPP4c together. 48 h post-transfection the damage was induced with X-ray irradiation (2 Gy). Microscopy images represent the foci (green) 8 h after X-ray irradiation in S phase (EdU, orange) cells. Scale bar: 10 µm. **D.** A total of 50 cells after RNAi (set #2 siRNA) were examined at various time points after irradiation to count the number of γH2Ax foci. The term "NT" refers to non-treated (non-irradiated) cells. The graphs display all data points along with the mean value and standard error of the mean (SEM) (n = 3). Asterisks on the graphs represent the P values obtained through linear regression, which were calculated independently for each time point. **E).** Validation of gene silencing by qPCR. 48 h post-transfection cells were harvested and the level of gene expression of silenced Ctb, PP4c, R3A or R3B with the second sets of siRNAs was measured by qPCR. The values were normalized to the level of actin.

**Supplementary Figure 3**

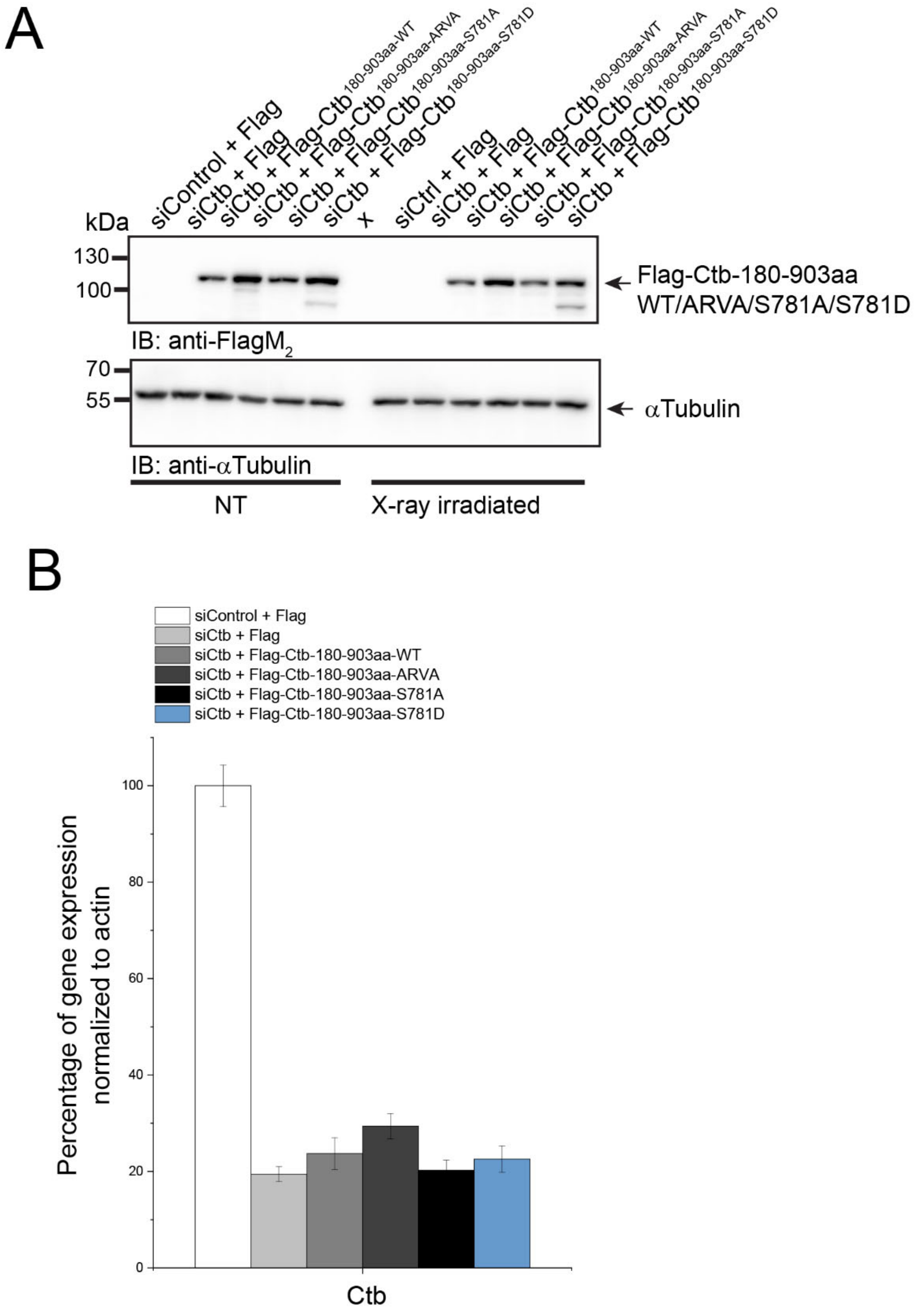


**Supplementary Figure 3. The FRVP and SQ motifs are crucial for Ctb-regulated DNA repair. A).** The expression level of wild-type or S781A/S781D/ARVA mutant variants of Flag-tagged $Ctb^{180\text{-}903aa}$ in control or Ctb-depleted cells was tested by Western blotting using the indicated antibodies. This shows comparable levels of the overexpressed Flag-tagged transgenic proteins. **B).** Validation of gene silencing by qPCR. 48 h post-transfection cells were harvested and the level of gene expression of silenced Ctb was measured by qPCR. The values were normalized to the level of actin.

**Supplementary Figure 4**

**Uncropped images corresponding to the main and supplementary figures (as indicated).**

Figure 1B (Uncropped)

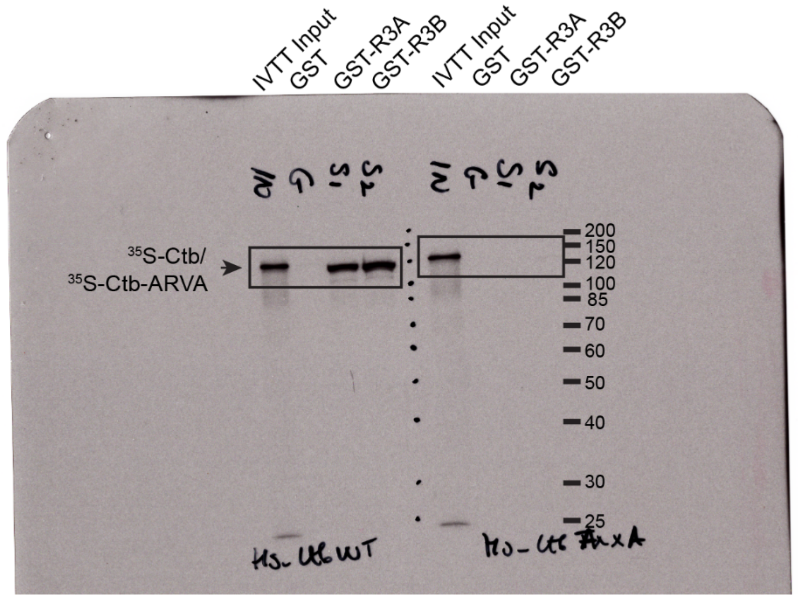

Figure 1C (Uncropped)

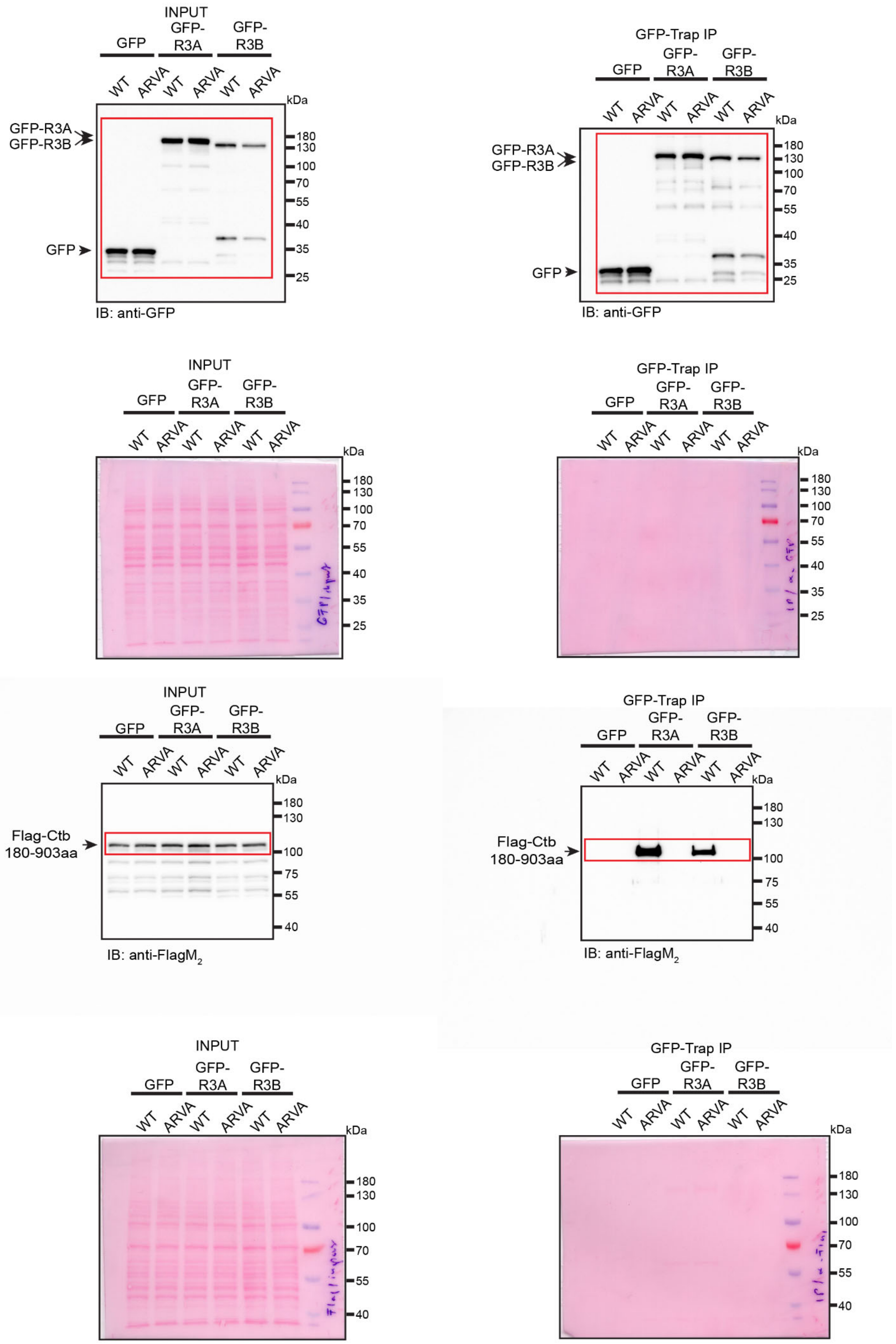

Figure 2D (Uncropped)

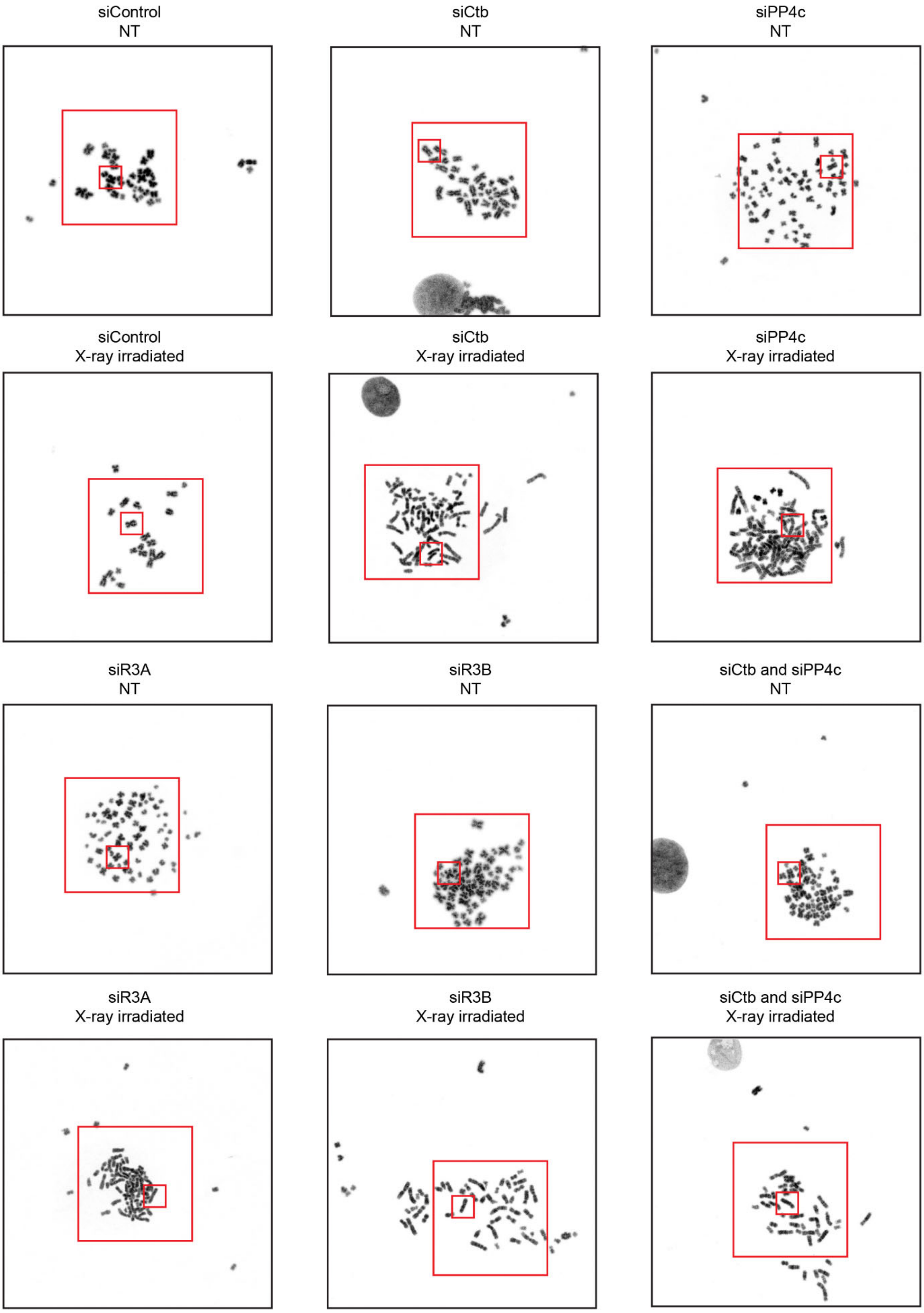

Figure 3A (Uncropped)

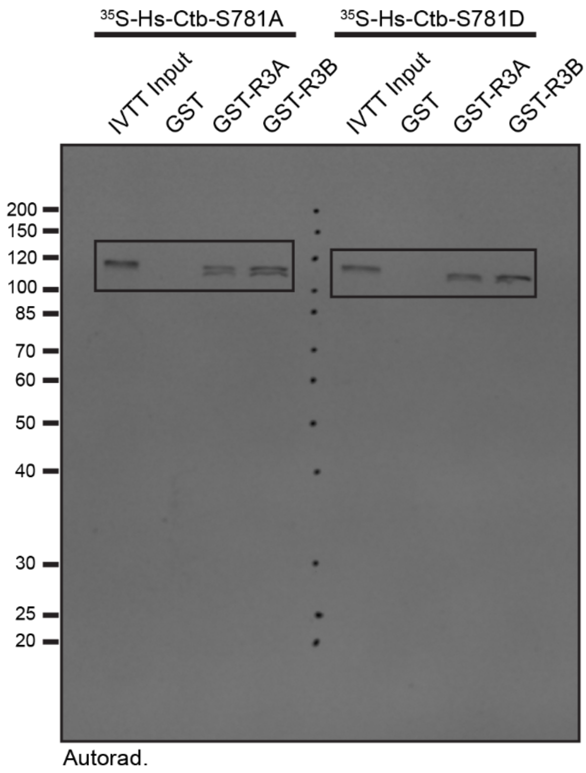


76

Figure 3B (Uncropped)

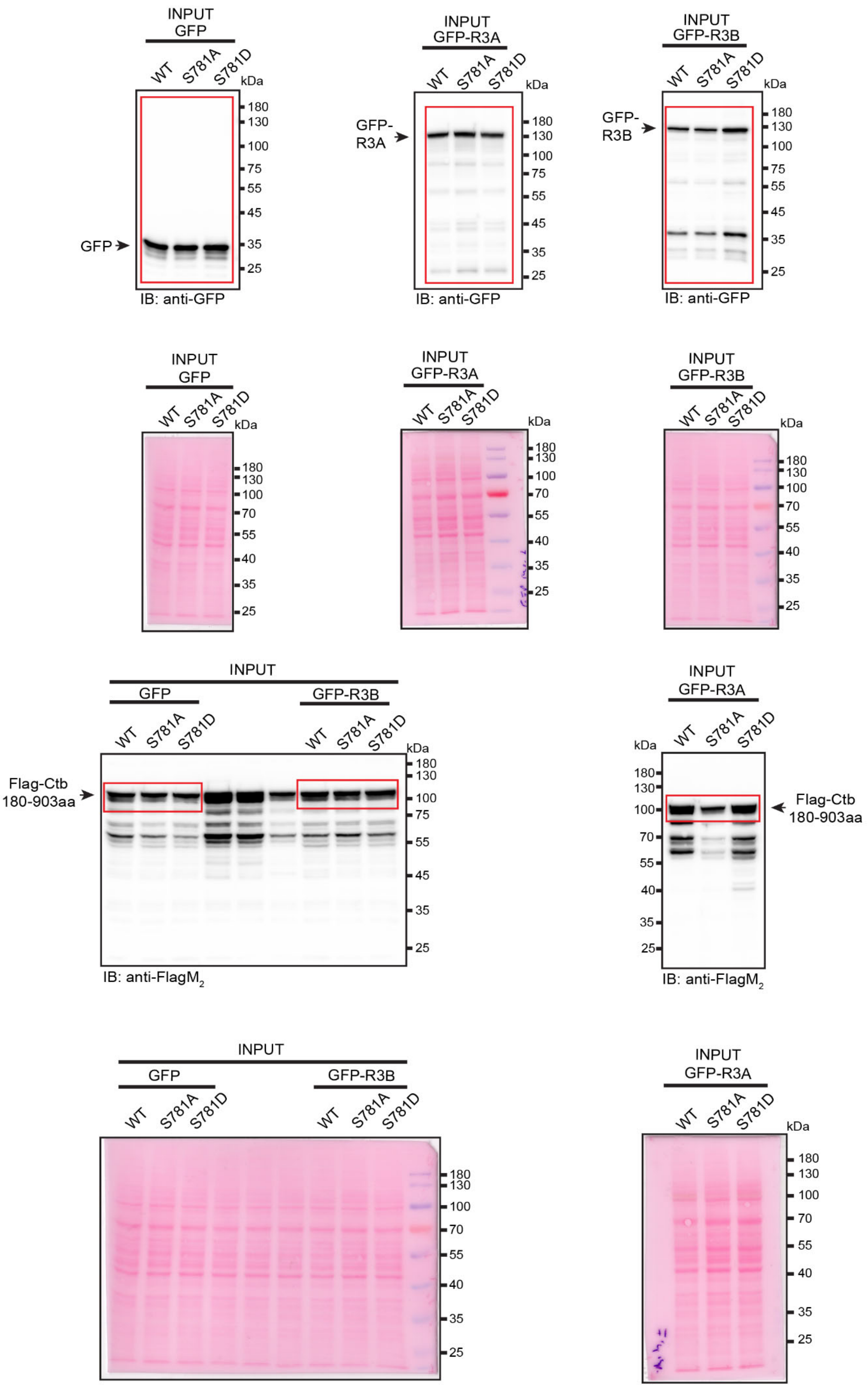

Figure 3B (Uncropped)

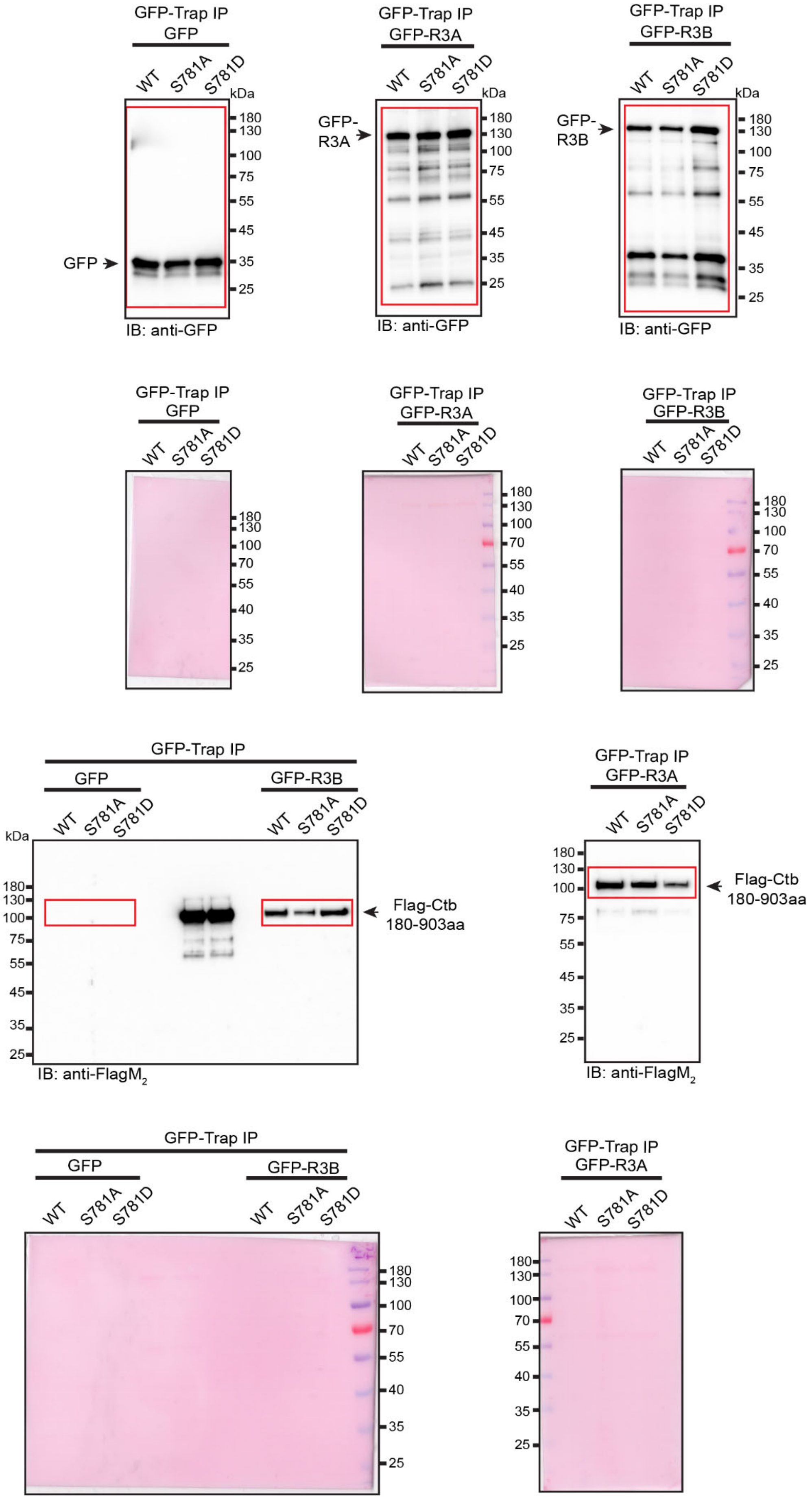

Figure 3D (Uncropped)

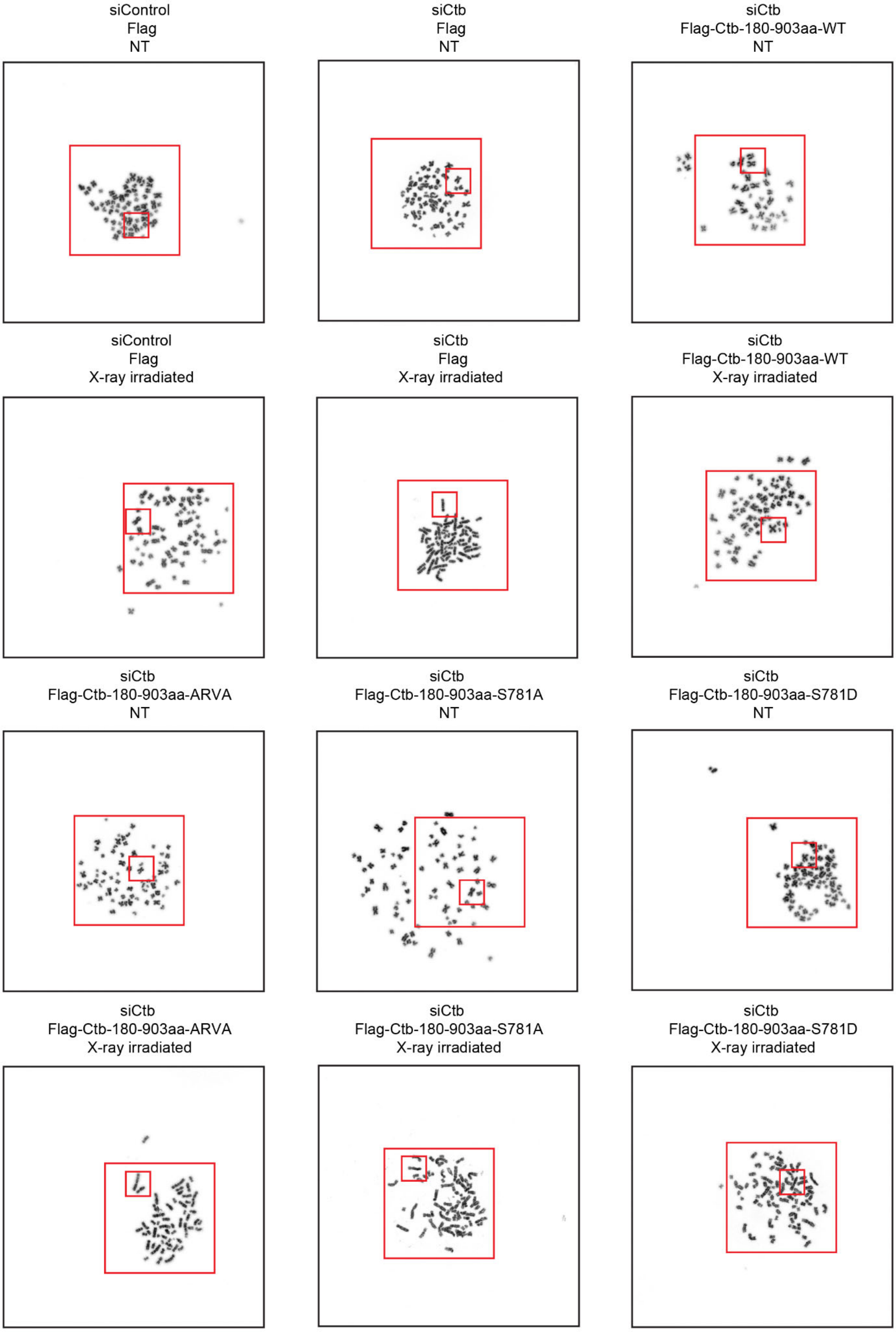


79

Supplementary figure 1B (Uncropped)

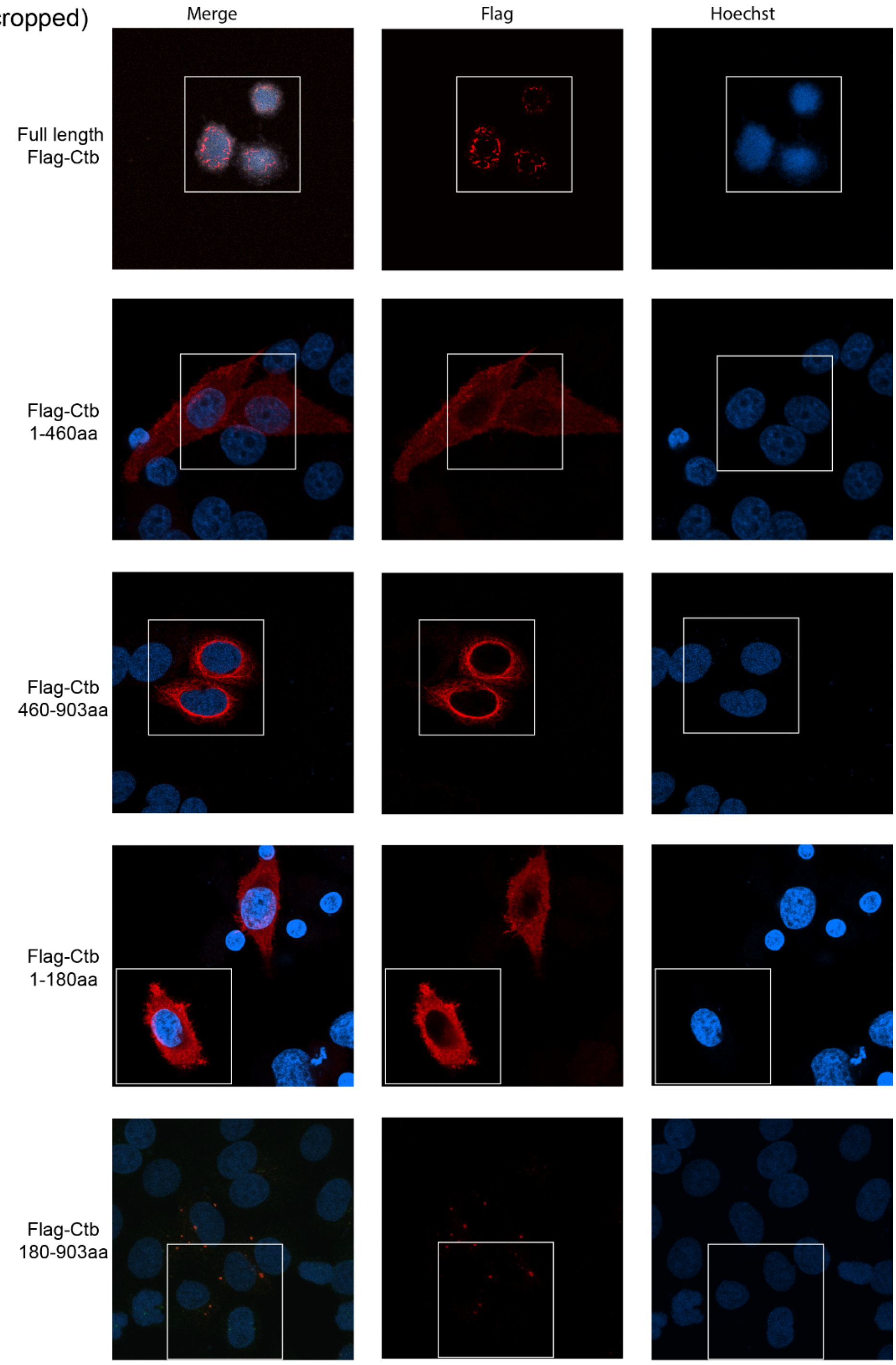


Supplementary figure 1C (Uncropped)

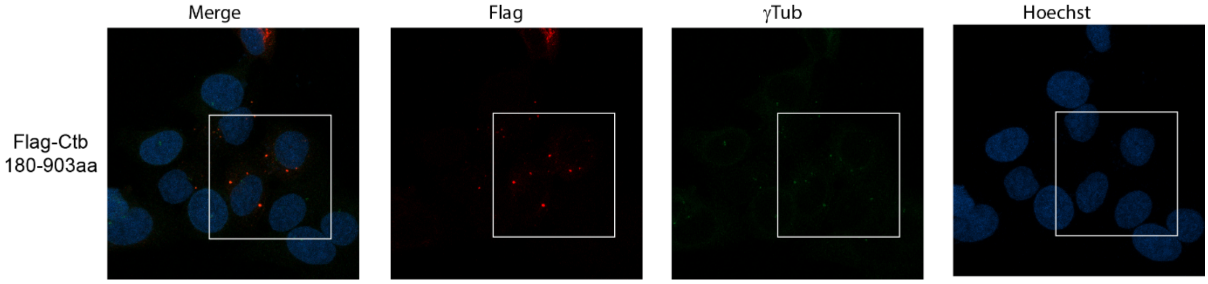

### Supplementary figure 3A (Uncropped)

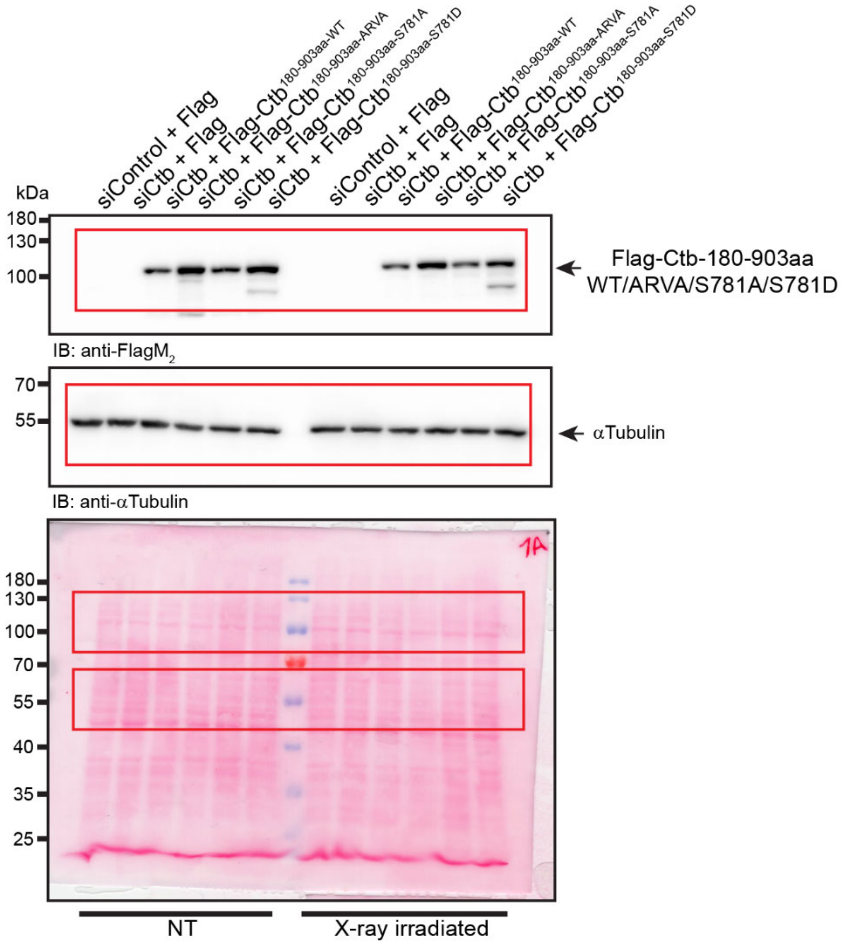


81
